# Respiratory heterogeneity shapes biofilm formation and host colonization in uropathogenic *Escherichia coli*

**DOI:** 10.1101/460311

**Authors:** Connor J. Beebout, Allison R. Eberly, Sabrina H. Werby, Seth A. Reasoner, John R. Brannon, Shuvro De, Madison J. Fitzgerald, Marissa M. Huggins, Douglass B. Clayton, Lynette Cegelski, Maria Hadjifrangiskou

**Affiliations:** Department of Pathology, Microbiology, and Immunology, Vanderbilt University Medical Center, Nashville, TN, USA; Department of Chemistry, Stanford University, Stanford, CA, USA; Division of Pediatric Urology, Vanderbilt University Medical Center, Nashville, TN, USA; Vanderbilt University, Nashville, TN, USA; Vanderbilt Institute for Infection, Immunology & Inflammation, Vanderbilt University Medical Center, Nashville, TN, USA

**Keywords:** biofilms, heterogeneity, oxygen gradients, respiration, urinary tract infection

## Abstract

Biofilms are multicellular bacterial communities encased in a self-secreted extracellular matrix comprised of polysaccharides, proteinaceous fibers, and DNA. Organization of these components lends spatial organization to the biofilm community such that biofilm residents can benefit from the production of common goods, while being protected from exogenous insults. Spatial organization is driven by the presence of chemical gradients, such as oxygen. Here we quantified and localized the expression of two *Escherichia coli* cytochrome oxidases in cells found in the biofilm state and defined their contribution to biofilm architecture. These studies elucidated a role for the high-affinity quinol oxidase cytochrome *bd* in matrix production and biofilm resident protection. Cytochrome *bd* was the most abundantly expressed respiratory complex in the biofilm community and was localized in the biofilm interior. Depletion of the cytochrome bd-expressing subpopulation led to decreased extracellular matrix and increased sensitivity of the community to exogenous stresses. Interrogation of the distribution of cytochrome oxidases in the planktonic state revealed that ∼15% of the population expresses cytochrome *bd* at atmospheric oxygen concentration, and this population dominates during acute urinary tract infection. These data point towards a bet-hedging mechanism in which heterogeneous expression of respiratory complexes ensures respiratory plasticity of *E. coli* across diverse host niches.

## Importance

Biofilms are multicellular bacterial communities encased in a self-secreted extracellular matrix comprised of polysaccharides, proteinaceous fibers, and DNA. Organization of these components lends spatial organization in the biofilm community. Here we demonstrate that oxygen gradients in uropathogenic *Escherichia coli* (UPEC) biofilms lead to spatially distinct expression programs for cytochrome oxidases - components of the terminal electron transport chain. Our studies reveal that the cytochrome *bd* expressing subpopulation is critical for biofilm development and matrix production. In addition, we show that cytochrome oxidases are also heterogeneously expressed in planktonic populations, and that this respiratory heterogeneity provides a fitness advantage during infection. These studies define the contributions of cytochrome oxidases to biofilm physiology and suggest the presence of respiratory bet-hedging behavior in UPEC.

## Introduction

Rather than existing as a phenotypically uniform population, bacterial communities are characterized by subpopulations that are oftentimes phenotypically distinct. This intra-strain heterogeneity can be transient, brought about by stochastic differences in the abundance and activity of regulators in each individual cell, or it can be irreversible, through the acquisition of mutations. Heterogeneity influences the fate of the particular strain by allowing for at least a portion of the population to thrive in different niches as a result of its specific expression program.

Biofilms are complex multicellular communities assembled in an organized fashion in three-dimensional space. One of the most critical features of biofilms is a self-secreted extracellular matrix (ECM) that comprises a variety of exopolysaccharides, proteinaceous fibers, and extracellular DNA (1). The ECM protects the biofilm residents from predation, desiccation, assault by antimicrobial agents, and, when biofilms form in the host, the immune system. In addition to providing a physical barrier against external threats, the ECM also serves as a barrier to diffusion. Restricted diffusion, in conjunction with the metabolic activity of resident bacteria, leads to the establishment of various chemical gradients throughout the biofilm community (2, 3). Bacteria at different locales along the gradient respond to the microenvironment differently, and as a result differentiate into distinct and often metabolically cooperative subpopulations (3–7). Previous studies in *Pseudomonas aeruginosa* and *Escherichia coli* indicated that oxygen gradients play a key role in regulating the differential expression of genes involved in biofilm formation and metabolic specialization (8–11). The presence of an oxygen gradient suggests the emergence of subpopulations that utilize different respiratory components as a function of the oxygen abundance to which they are exposed. This leads to the hypothesis that the metabolic program of differentially respiring subpopulations is distinct from one another and contributes to differential production of biofilm goods that in turn enhance biofilm resilience.

*Escherichia coli* is a facultative anaerobe capable of utilizing multiple metabolic pathways to fulfill its energy requirements. In aerobically respiring *E. coli*, cytochrome oxidases comprise essential components of the terminal electron transport chain that couple the flow of electrons to the reduction of molecular oxygen into water (12, 13). *E. coli* encodes two classes of cytochrome oxidases with differing oxygen affinities: one heme copper oxidase, cytochrome *bo* (encoded by the *cyoABCD* gene cluster), and two bd-type oxidases, cytochromes *bd (cydABX)* and *bd*_2_ *(appBC)* (13, 14). Cytochrome *bo* is induced at high (atmospheric, 21%) oxygen tensions, whereas the bd-type oxidases are induced at low (hypoxic, 2-15%) oxygen tensions (13, 15). Based on these *in vitro* expression patterns, we hypothesized that cytochrome *bo* would be enriched on the air-exposed biofilm surface, whereas cytochromes *bd* and *bd*2 would be enriched in the hypoxic interior.

Here we report that the spatial distribution of cytochrome oxidases in biofilms formed by uropathogenic *Escherichia coli* (UPEC) is a fundamental driver of biofilm architecture. Peptide nucleic acid fluorescence *in situ* hybridization (PNA-FISH) analyses assigned locations to each cytochrome-producing subpopulation, elucidating for the first time spatially distinct expression programs for respiratory oxidases in *E. coli*. Depletion of the cytochrome *bd*-expressing subpopulation from the biofilm significantly impaired diffusion resistance by altering the abundance and organization of the ECM. Assessment of deletion mutants in a well-established urinary tract infection murine model revealed that only the cytochrome *bd* mutant was significantly attenuated for virulence, although the infecting pool of bacteria in the parent strain exhibited heterogeneous expression of all three respiratory oxidases. *In situ* analysis of urine-associated bacteria demonstrated a shift of the population to cytochrome *bd* expression, suggesting that the bladder favors cytochrome *bd*-expressing bacteria and that heterogeneity in the input pool provides a fitness advantage to uropathogenic strains. Our studies, performed on one of the most commonly acquired human pathogens and a prolific biofilm producer *in vivo*, unveil a potential avenue for targeting heterogeneity and homogenizing bacterial programming as a therapeutic approach.

## Results

### Cytochrome bd is the most abundant respiratory transcript in mature UPEC biofilms

To test the hypothesis that multiple differentially respiring subpopulations exist in *E. coli* biofilms, we measured the relative expression of components of the aerobic *(cyoABCD, appBC*, and *cydABX)* and anaerobic *(napABC* and *narGHI)* respiratory chains in mature colony biofilms. Previous studies demonstrated that the central and peripheral regions of biofilms are metabolically distinct, yet co-dependent (16). The central region is shielded from environmental stressors and is therefore critical for survival of the population. The peripheral region contains a rapidly expanding population that contributes most to biofilm growth at the expense of being highly susceptible to exogenous threats. We first analyzed transcript levels in the center and periphery of colony biofilms grown on Yeast Extract/Casamino Acids (YESCA) agar (Fig. 1a), a medium which induces expression of key matrix components (curli amyloid fibers, type 1 pili, secreted proteins Hu-α/β, cellulose) that are also critical for fitness in the urinary tract (10, 17–19). At both regions, of the five respiratory operons analyzed, over 90 percent of transcripts corresponded to aerobic respiratory components (Fig. 1b-c). The most abundant transcript was that of *cydA* (Fig. 1b-d), corresponding to cytochrome *bd* complex *(cydABX)* expression. The *cydA* transcript levels were approximately 2-fold higher than those corresponding to *cyoABCD* (Fig. 1b-d), which was the second most highly abundant oxidase under the conditions tested. Cytochrome *bd*_2_ was the third most-highly expressed, while the anaerobic *narG* and *napA* genes exhibited baseline expression levels (Fig. 1b-d). These results validated that most of the biofilm is hypoxic, consistent with previous work in *P. aeruginosa* and *E. coli* (8, 9).

**Figure 1.**
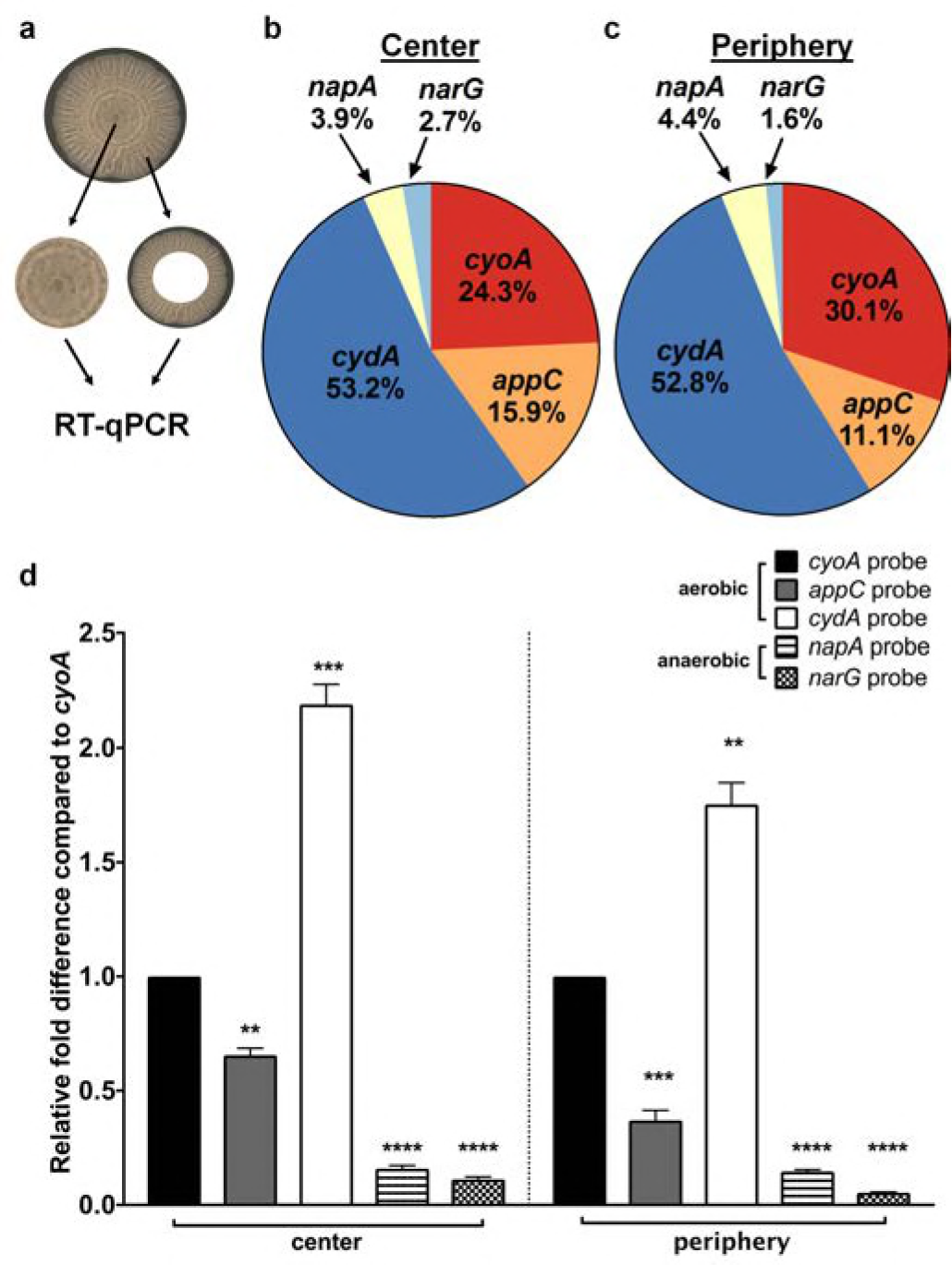
Lateral expression of respiratory complexes in *Escherichia coli* biofilms: **a,** Image of a mature colony biofilm formed by UPEC strain UTI89 on YESCA agar. The center and periphery of colony biofilms were harvested and subjected to RNA extraction and RT-qPCR using the following probes: *cydA, cyoA, appC, napA*, and *narG* with *gyrB* as a normalizer. **b – c**, Pie charts indicating the relative abundance of detected respiratory transcripts in the biofilm center (**b**) and periphery (**c**). **d,** Graphs depicting relative fold differences in transcript abundance in the biofilm center and periphery. The graph and pie chart depict the average of 4 biological replicates. Statistical analysis was performed in GraphPad Prism using a two-tailed paired t test. Data are presented as mean ± SEM. * = p < 0.05, ** = p < 0.01, *** = p< 0.001, **** = p < 0.0001.

### Spatial organization of cytochrome expression along the oxygen gradient

Previous studies elucidated the presence of steep biofilm oxygen gradients as a function of biofilm depth (Fig. 2a) (8, 9). To define the spatial distribution of cytochrome-expressing subpopulations, we performed PNA-FISH on cryosections of mature colony biofilms using probes targeting each cytochrome oxidase operon (cyoA, *appC*, and *cydA)* as well as *rrsH* as an endogenous control. Each PNA-FISH probe was designed using the validated probe sequences used for qPCR (Fig. 1 and Table S1) to ensure comparable hybridization efficiencies for each probe. SYTO 9 staining of sections was used as an additional control to localize the entire biofilm community and account for possible hybridization inconsistencies with the *rrsH* control probe (Figs. 2d, j and S1). Consistent with previous observations demonstrating that the highest oxygen abundance is at the air-exposed surface of the biomass (8, 9), we observed that *cyoABCD* transcript was most abundant in bacteria lining the air-exposed surface of the biofilm (Fig. 2b-c, f, h-i, l, n). In contrast, the highest abundance of *cydABX* transcript was found in densely packed clusters of bacteria in the interior of the biofilm (Fig. 2b-c, e, g, i, k, m). Interestingly, while in K12 *E. coli* the function of cytochrome *bd* and *bd_2_* are thought to be redundant, the expression distribution of the two gene clusters was different, with *appBC* observed to be evenly distributed throughout the community (Fig. S1).

**Figure 2.**
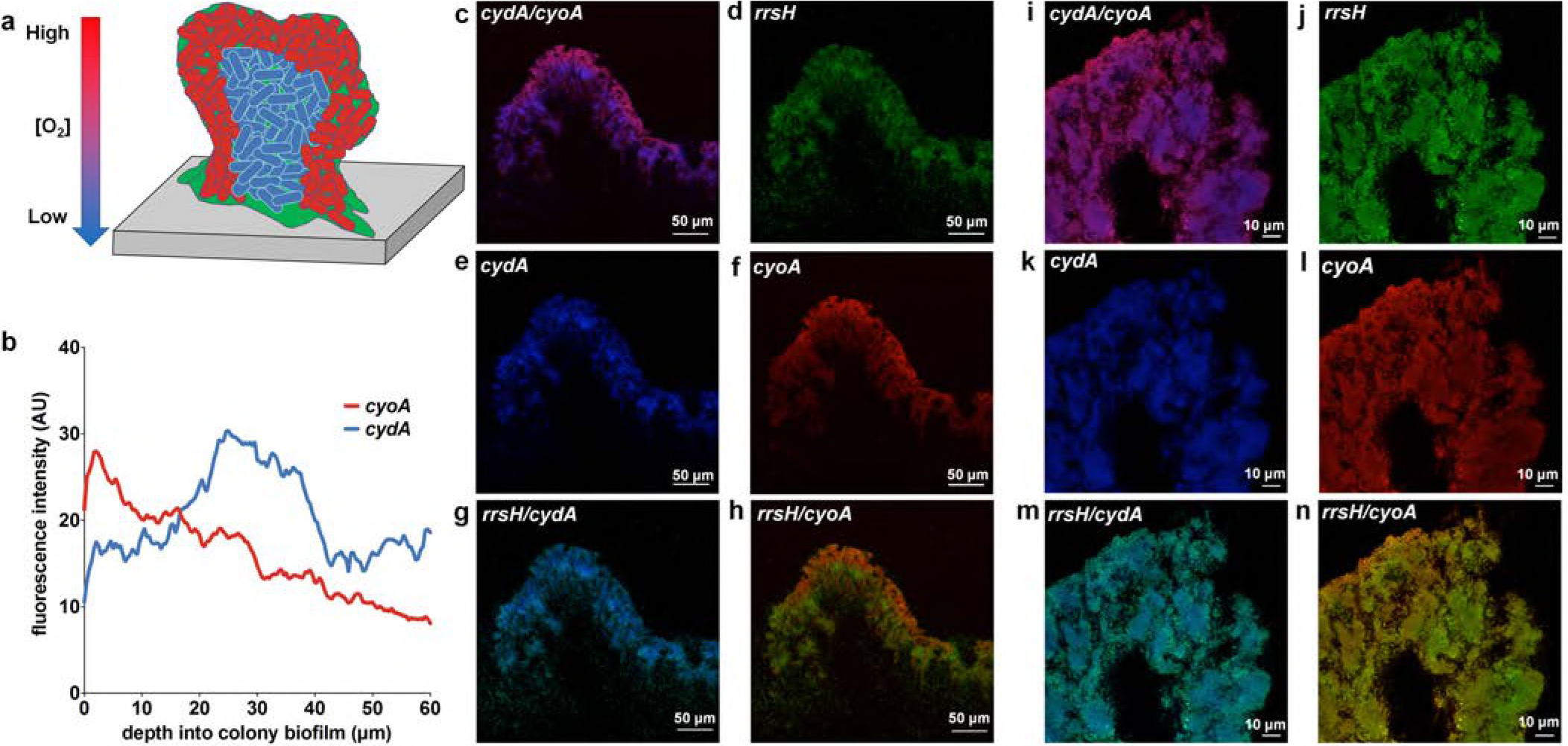
Expression of cytochrome oxidases as a function of the oxygen gradient. **a**, Cartoon depicting the detected localization of *cyoA* (red) and *cydA* (blue) transcripts in biofilm cryosections. **b**, Fluorescence intensity of *cyoA* and *cydA* PNA-FISH probes were quantified on ImageJ. Data are presented as the average fluorescence intensity as a function of depth obtained from four images, each with five measurements per image. **c – n**, Representative images of PNA-FISH stained biofilm cryosections at 20x magnification (**c – h**) and 63x magnification (**i – n**). Cryosections were stained with PNA-FISH probes targeting *cyoA* and *cydA* with *rrsH* (16S rRNA) as an endogenous control. Images are representative of three biological replicates.

### Loss of cytochrome bd alters biofilm architecture, development, and matrix abundance

The highly regulated spatial expression of cytochrome *bo* and cytochrome *bd* oxidases in the biofilm raised the hypothesis that each of these cytochrome-expressing subpopulations uniquely contributes to overall biofilm architecture. To test this hypothesis, we created isogenic deletion mutants lacking the *cyoAB, appBC*, or *cydAB* genes and compared the biofilms formed by the resulting strains (Fig. 3a). Colony biofilms formed by the parental strain expand to an average diameter of 16.8 mm over an 11-day incubation period and exhibit elaborate rugose architecture with distinct central and peripheral regions (Figs. 3a and S2). Strains deleted for *cyoAB* and *appBC* exhibited inverse phenotypes to each other, with the *ΔcyoAB* colony biofilms expanding more than the parental strain (average diameter: 19.9 mm) and the *ΔappBC* colony biofilms appearing more compact and with apparently higher rugosity (Figs. 3a and S2). Strikingly, while *ΔcyoAB* and *ΔappBC* only displayed minor architectural changes, *ΔcydAB* colony biofilms exhibited pronounced defects both in development and architecture (Fig. 3a and S2). Colonies deleted for *cydAB* grew at a similar rate as the parent, *ΔcyoAB*, and *ΔappBC* for the first 72 hours (Fig. S2). However, colony growth was significantly stunted between days 3 and 11, with radial expansion remaining at an average diameter of 10.3 mm and colonies exhibiting a wet mass of 48.11 mg (70% of the parent strain) at the end of the experiment, even though the CFU produced by the two strains were comparable across the 11-day incubation period (Figs. 3b-c and S2). Complementation of *ΔcydAB* with an extra-chromosomal construct expressing *cydABX* under its native promoter rescued the deletion phenotype, indicating that the defects observed in the *ΔcydAB* mutant stem solely from the removal of the *cydABX* cluster (Fig. S3). Together, these observations demonstrate that cytochrome *bd* is a key contributor to biofilm development and suggest that loss of *cydAB* alters the synthesis and organization of the ECM.

**Figure 3.**
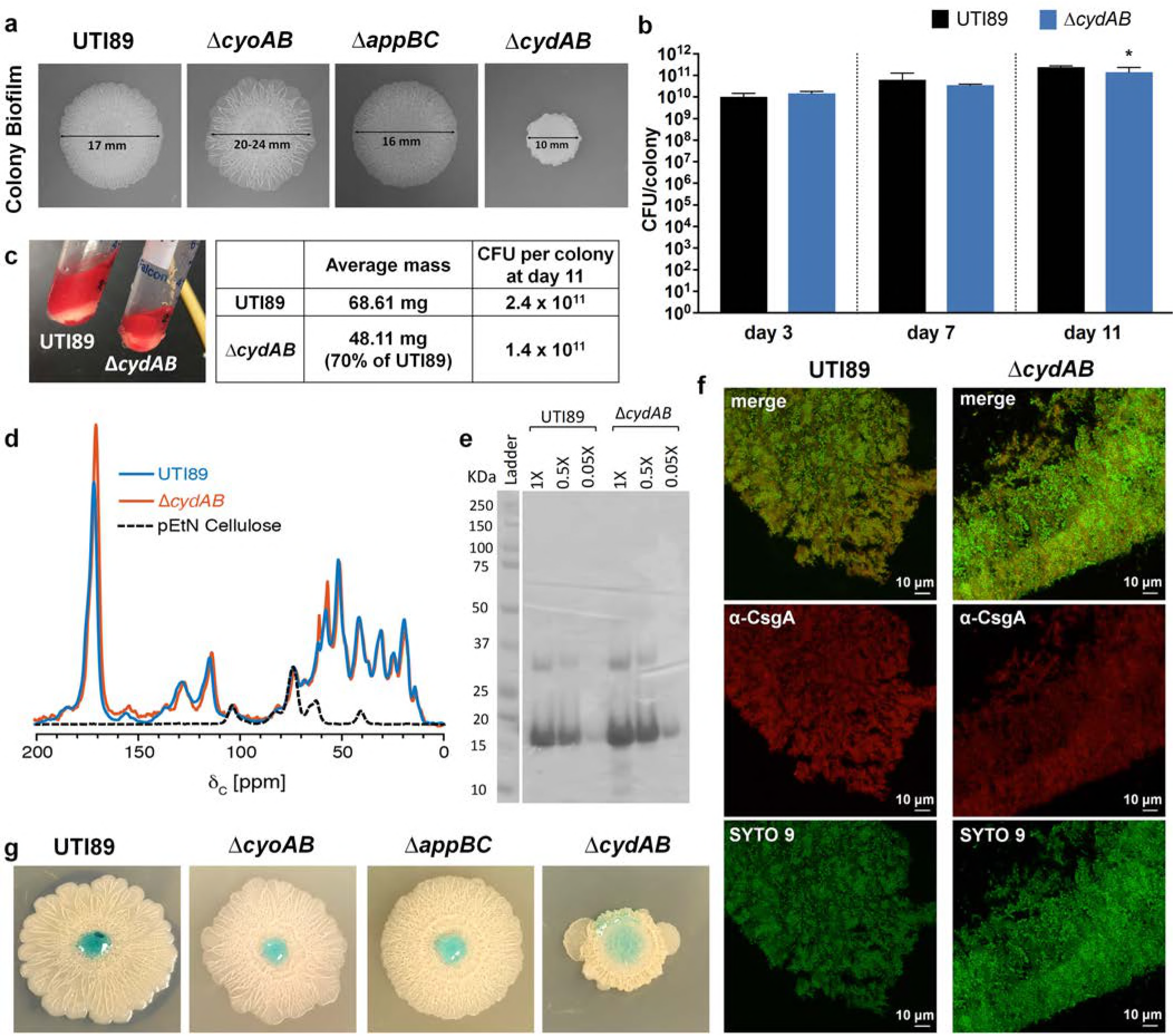
Cytochrome *bd* organizes biofilm architecture and ECM production. **a,** Colony biofilms of UTI89 and cytochrome oxidase mutants grown on YESCA agar for 11 days. Images are representative of at least 30 biological replicates. **b**, CFU per colony biofilm was measured at days 3, 7, and 11 of growth. Statistical analysis was performed in GraphPad Prism using a two-tailed unpaired t test. * = p < 0.05. Data are presented as mean ± SD. Data are representative of five biological replicates. **c, left**, Image depicting gross changes to extracellular matrix (ECM) abundance between UTI89 and *ΔcydAB* colony biofilms. ECM is stained red by the presence of Congo Red in the growth medium. **right**, Table lists average mass and CFU per colony biofilm after 11 days of growth. Average mass is from a pooled analysis of 186 individual colonies. CFU per colony is the average of five biological replicates. **d**, Solid-state NMR spectra of the ECM of UTI89 (blue), *ΔcydAB* (orange), and isolated pEtN cellulose (black). **e**, SDS-PAGE gel of UTI89 and *ΔcydAB* ECM. **f**, Immunofluorescence images of curli (α-CsgA, red) localization in UTI89 and *ΔcydAB* colony biofilm cryosections. **g**, Colored water droplets were added to the top of day 11 colony biofilms to probe biofilm barrier function.

Under the conditions used, the ECM of *E. coli* comprises primarily of cellulose and curli amyloid fibers (20). Previous solid-state nuclear magnetic resonance (NMR) spectroscopy analyses on intact ECM material defined the contributions of cellulose and curli to the *E. coli* biofilm ECM, and determined that curli and cellulose are present in a 6 to 1 ratio (20). More recently, the ECM cellulose was determined to be a chemically modified form of cellulose, specifically phosphoethanolamine (pEtN) cellulose (21). To interrogate the effects of *cydAB* deletion on curli and exopolysaccharide production, we extracted ECM and performed solid-state NMR analysis to evaluate the abundance of curli and cellulose components (Fig. 3d-e). The NMR spectra obtained for the parent and *ΔcydAB* ECM are very similar overall, indicating a comparable protein to polysaccharide ratio between the samples (Fig. 3d). Consistent with this analysis, we observe no change in protein composition between the parent and *ΔcydAB* ECM samples when analyzed on SDS-PAGE gels (Fig. 3e). We additionally do not observe any overt alterations to curli abundance or localization between UTI89 and *ΔcydAB* biofilm cryosections using immunofluorescence (Fig. 3f). Despite the similar composition, the total amount of ECM recovered was reduced in *ΔcydAB*, indicative of a decrease in ECM production. Because the protein to polysaccharide ratio and curli abundance are unchanged between the parent and *ΔcydAB*, these data are suggestive of a change to the overall mixture of matrix components in *ΔcydAB*, with particular reductions in the abundance of non-curli and non-pEtN cellulose ECM components.

The ECM plays a central role in biofilm physiology by providing physical protection against exogenous insults, serving as a structural scaffold, and helping to establish chemical gradients which lead to metabolic differentiation and subpopulation formation (2, 3, 17, 22). As such, disruptions to the matrix can have catastrophic consequences for the biofilm community. We hypothesized that the altered matrix abundance and architecture in the *ΔcydAB* mutant would render the biofilm more susceptible to exogenous insults. To investigate this possibility, we probed the barrier function of the cytochrome oxidase mutant biofilms by applying a drop of colored water to the surface of mature colony biofilms (Fig. 3g and Movies S1-4) (23). While the parent strain, *ΔcyoAB*, and *ΔappBC* biofilms repelled the drop, the solution readily penetrated *ΔcydAB* biofilms, demonstrating that the alterations to *ΔcydAB* biofilm architecture and matrix abundance increases penetrance of aqueous solutions.

### Loss of cytochrome bd increases population sensitivity to nitrosative stress under ambient oxygen concentrations

Together, our studies indicate that cytochrome *bd* is highly expressed in biofilms, and that loss of the cytochrome bd-expressing subpopulation impairs barrier function and reduces the abundance of extracellular matrix. These data suggest that the cytochrome bd-expressing subpopulation plays a critical role in promoting ECM synthesis and providing structural integrity to the community. However, it is also possible that cytochrome *bd* is preferentially expressed in the biofilm because cytochrome *bd* provides protection against oxidative and nitrosative stress - byproducts of biofilm metabolism (24, 25) and, in the case of infection, components of the innate immune response (26–28). Previous studies demonstrated that cytochrome *bd* has catalase activity, is capable of oxidizing the respiratory inhibitor nitric oxide, and is insensitive to nitrosative stress due to its unusually fast nitric oxide dissociation rate (26). These biochemical activities are thought to occur at unique locations on the protein; quinol oxidation occurs at the periplasmic Q loop, oxygen reduction and nitric oxide binding occurs at heme d, and catalase activity is thought to occur through heme b_595_ (14, 29, 30). By contrast, cytochrome *bo* affords no protection against oxidative or nitrosative stress and is irreversibly inhibited by nitric oxide (26). Given these additional functions of cytochrome *bd*, we performed growth curves at ambient oxygen concentration and evaluated the effects of nitrosative and oxidative stress on the fitness of cells lacking *cydAB* or *cyoAB* as compared to the parent strain. Without the addition of stress, both mutants exhibited a lag in growth (Fig. 4a), but each reached a maximal CFU/mL similar to the parental strain by the end of the growth experiment (Fig. 4a). ATP measurements of the corresponding cultures revealed no significant overall differences in ATP concentrations between the strains during logarithmic growth (Fig. 4b). Next, to determine whether loss of cytochrome *bd* impairs resistance to oxidative and nitrosative stress, we measured growth with and without these stressors. Whereas treatment with H_2_O_2_ had no significant effects on the growth of the two mutants (Fig. 4c), addition of the nitric oxide donor NOC-12 to planktonic cultures most significantly impacted growth of *ΔcydAB* (Fig. 4d), consistent with previous studies on K12 *E. coli* and the multi-drug resistant strain ST131 (27, 28). Together, these data demonstrate that while cytochrome *bd* is dispensable for energy generation during planktonic growth, loss of cytochrome *bd* sensitizes bacteria to nitrosative stress. In conjunction with the disrupted biofilm architecture and altered matrix abundance in *ΔcydAB* biofilms, these data suggest that cytochrome bd-expressing subpopulations are critical, not only for directing matrix biosynthesis, but also for withstanding harmful metabolic byproducts while in the biofilm state.

**Figure 4.**
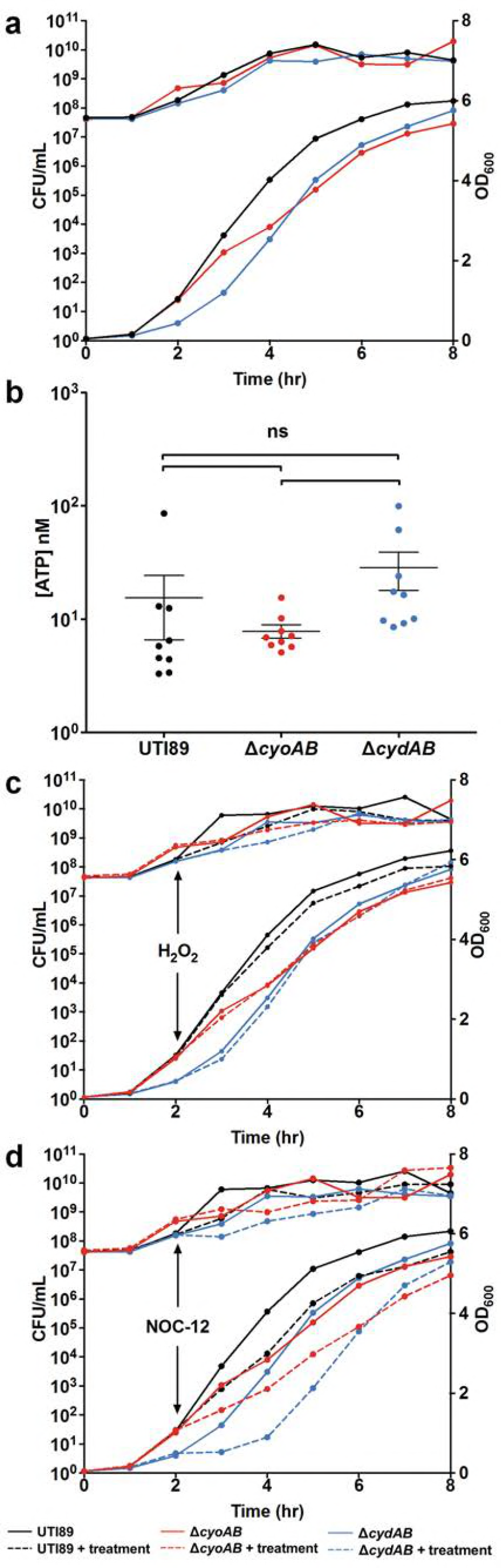
Cytochrome *bd* provides nitrosative stress resistance. **a**, Growth curves for UTI89, *ΔcyoAB*, and *ΔcydAB* as measured by CFU per mL (upper lines, left axis) and OD_600_ (lower lines, right axis). **b**, ATP levels measured from logarithmic cultures of UTI89, *ΔcyoAB*, and *ΔcydAB*. Statistical analysis was performed on GraphPad Prism using a two-tailed unpaired t test. Data are presented as mean ± SEM. **c – d**, Growth curves for UTI89, *ΔcyoAB*, and *ΔcydAB* as measured by CFU per mL (upper lines, left axis) and OD_600_ (lower lines, right axis). 0.5 mM of nitric oxide donor NOC-12 (**c**) or 1 mM hydrogen peroxide (**d**) was added at t = 2 hours. All data are representative of at least three biological replicates.

### Heterogeneous expression of cytochrome oxidases at the population level

Our data thus far indicate that in addition to heterogeneity in cytochrome oxidase expression in the biofilm state, heterogeneous expression of cytochrome oxidases must also be occurring in the planktonic population. Our planktonic studies revealed a lag in growth of the *ΔcyoAB* and the *ΔcydAB* mutants when these strains were grown under ambient oxygen concentrations, suggesting that in a given culture there are subpopulations - like in the biofilm - that stochastically or deterministically express different respiratory components. Such a bet-hedging approach could provide UPEC with the flexibility to quickly adapt to a given niche, be it different locales in the genitourinary tract or in the gastrointestinal tract during host colonization. In the context of urinary tract infection, *E. coli* traverse from the nearly anoxic gut to the perineum, where it encounters atmospheric oxygen concentrations, prior to ascending the urethra to enter the hypoxic bladder, where the dissolved urinary oxygen concentration is 4 – 6% (31). This microbial journey is performed by planktonic cells, which can then expand into multicellular communities on and within bladder epithelial cells, as well as on urinary catheters (1, 32). In previous studies, we and others demonstrated that UPEC respire aerobically during infection (33–35) and that biofilm formation is favored in conditions that mimic oxygen levels in the bladder (36).

The high abundance of *cydABX* transcript in the hypoxic areas of the biofilm, in conjunction with the defects observed in aerobically grown *ΔcydAB* planktonic cultures, raised the hypothesis that a cytochrome bd-expressing subpopulation exists in the planktonic state under ambient oxygen conditions and that this cytochrome *bd* expressing subpopulation exhibits the greatest fitness advantage during infection. To test this hypothesis, we first analyzed transcript abundance in cultures used for inoculation during murine infections with qPCR and PNA-FISH (Fig. 5). Under these conditions, qPCR analysis indicated that the majority of transcript corresponds to *cyoABCD* (69.7%), with *cydABX* and *appBC* transcripts each comprising approximately 15% of detected transcripts (Fig. 5a). Transcript abundance was altered by decreasing ambient oxygen concentrations, with the most abundant transcript corresponding to *cydABX* in 12%, 8% and 4% oxygen, the latter being the concentration of dissolved oxygen concentration in the urine (Figs. 5a-b and S4) (31). PNA-FISH analysis revealed the presence of bacteria which uniquely express cytochrome *bo* (Fig. 5f), *bd* (Fig. 5e), or *bd_2_* (Fig. 5g), as well as some cells that have transcript of all three operons (Fig. 5c, e-g). Intriguingly, we observed dividing cells in which each daughter had distinct cytochrome oxidase transcript abundance (Fig. 5c, inset), suggesting that asymmetric distribution of respiratory transcripts during division may be a mechanism by which these subpopulations are generated.

**Figure 5.**
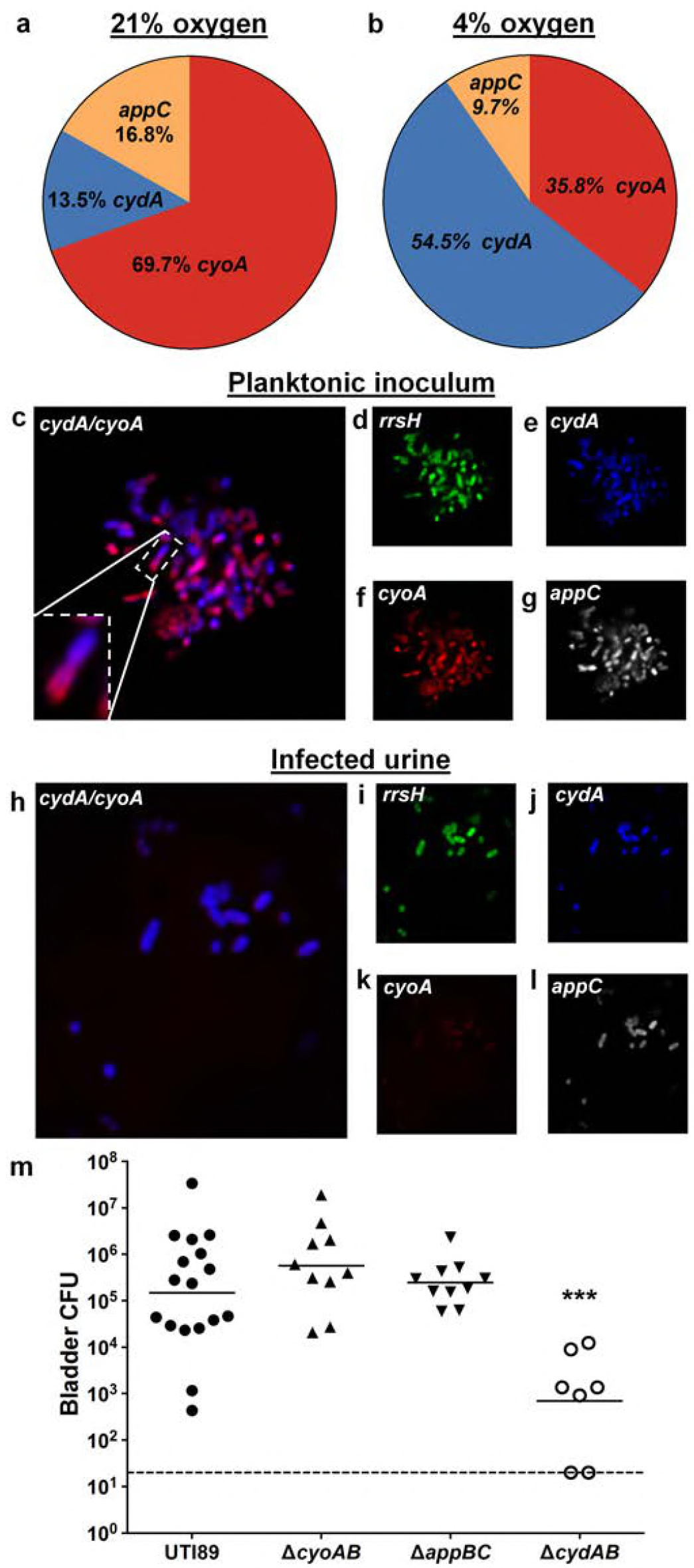
Respiratory heterogeneity provides afitness advantage during urinary tract infection. **a – b**, Pie charts depicting relative abundance of *cydA,cyoA*, and *appC* transcripts detected using RT-qPCR in planktonic cultures grown at 21% oxygen in the manner used to prepare cultures to inoculate mice (**a**), as well as planktonic cultures grown at 4% oxygen (**b**). Data is representative of three biological replicates. **c – g**, PNA-FISH was used to detect cytochrome oxidase transcripts from cultures used to inoculate mice. Data is representative of three biological replicates. **h – l**, PNA-FISH was used to detected cytochrome oxidase transcripts in the urine of mice infected with UTI89. Urine was pooled from 20 mice. **m**, Graph depicting bladder titers obtained from mice infected with UTI89 or cytochrome oxidase mutant strains at 24 hours post infection. Each point represents a mouse. UTI89 and *ΔcydAB* are representative of two independent experiments. *ΔcyoAB* and *ΔappBC* are representative of one experiment. Statistical analysis was performed in GraphPad Prism using a two-tailed Mann-Whitney test. Line represents geometric mean. *** = p<0.001.

### Expression of cytochrome bd is dominant during acute urinary tract infection

Previous studies reported that deletion of cytochrome *bd* impairs UPEC virulence in a UTI model (28). To gauge the contribution of each cytochrome oxidase during infection, we evaluated the fitness of *ΔcyoAB, ΔappBC*, and *ΔcydAB* mutants compared to the parent strain in a murine model of acute urinary tract infection. Consistent with the previous report (28), *ΔcydAB* exhibited a ∼2 log decrease in bladder colonization by 24 hours relative to the parent strain, while the mutants deleted for *cyoAB* and *appBC* colonize mice at the same level as the parent strain (Fig. 5m). Subsequent PNA-FISH on pooled urine obtained from mice infected with the parent strain revealed a marked enrichment in cytochrome bd-expressing cells and a corresponding reduction in the number of cells expressing cytochrome *bo* (Fig. 5h-l). This suggests that the bladder environment either induces transcription of *cydABX* or that only subpopulations of bacteria expressing *cydABX* are capable of efficiently colonizing the bladder. Together these data reveal the presence of subpopulations of bacteria that differentially express cytochrome oxidases as a potential bet-hedging mechanism to promote bladder colonization.

## Discussion

Cytochrome *bd* is a multifunctional protein that is central to respiration and can maintain activity in the face of nitrosative stress (26). As such, bacteria expressing cytochrome *bd* presumably exhibit a fitness advantage during growth conditions that are low in oxygen or high in metabolic byproducts that increase nitric oxide concentration. The biofilm state, while protecting the bacterial residents from predation and desiccation, constitutes a high-density environment with several chemical gradients that result from the consumption and production of metabolites. Accordingly, expressing an enzyme that can facilitate tolerance to metabolic byproducts, such as nitric oxide, would ensure that biofilm residents do not perish as a consequence of their own metabolic excretions. Our study elucidates the distribution of cytochrome oxidase expression in the biofilm state and indicates that the bulk of biofilm residents express cytochrome *bd*, particularly in the densely populated interior. The cytochrome bd-expressing bacteria are not necessarily using cytochrome *bd* for respiration, as many of them also have low levels of cytochrome *bo* and *bd_2_* transcripts (Fig. 2 and S1). Rather, the production of cytochrome *bd* may be leveraged towards providing tolerance to nitrosative stress, which irreversibly inhibits cytochrome *bo.* Indeed, in *ΔcydAB* biofilms we observe a marked increase in cytochrome *bo* expression (Fig. S5), suggesting that loss of cytochrome *bd* impairs nitric oxide tolerance and that increased production of cytochrome *bo* may be a compensatory mechanism that allows biofilm bacteria to respire in the presence of high levels of nitric oxide.

In addition to acting as a respiratory inhibitor, nitric oxide regulates cyclic di-GMP abundance and thereby governs the switch from motility to aggregation and biofilm expansion (37, 38). Consequently, if cytochrome *bd* decreases nitric oxide availability, it would indirectly influence ECM production. Consistent with this hypothesis, loss of the cytochrome bd-expressing subpopulation reduces the total abundance of matrix components and leads to gross alterations of biofilm architecture (Fig. 3). It is thus possible that the cytochrome bd-expressing subpopulation is critical for promoting the biosynthesis of the ECM by influencing the nitric oxide - cyclic di-GMP signaling axis. We are currently investigating this possibility.

Most importantly, this work revealed the presence of planktonic subpopulations that express distinct cytochrome oxidases during growth. In conjunction with the observation that only cytochrome *bd* expression is critical for fitness during infection, this finding suggests that basal expression of cytochrome *bd* under aerobic conditions serves as a bet-hedging mechanism that promotes the expansion of bacteria during the transition from the aerobic perineum to the hypoxic bladder. In addition to allowing for efficient respiration in the hypoxic bladder, expression of cytochrome *bd* provides resistance against nitrosative stress - a metabolic byproduct and component of the innate immune response - and promotes the formation of resilient biofilm communities. These observations suggest the presence of respiratory bet-hedging behavior in UPEC, and additionally suggest the possibility of targeting heterogeneity as a method for homogenizing bacterial populations and impeding their ability to colonize the urinary tract.

## Materials and Methods

### Bacterial Strains

All studies were performed in *Escherichia coli* cystitis isolate UTI89 (39). All gene deletions (*ΔcyoAB, ΔappBC*, and *ΔcydAB*) were performed using the λ-red recombinase system (40). Complementation constructs were created in plasmid pTRC99a with *cydABX* under the control of its native promoter as previously described (41). Primers used for gene deletions and complementation plasmid construction are listed in Table S1.

### Growth conditions

For all analyses, strains were propagated overnight at 37°C with shaking in Lysogeny broth (LB) (Fisher) at pH 7.4. To form colony biofilms, 10 μL of overnight culture was spotted onto 1.2x Yeast Extract/Casamino Acids (YESCA) agar (9) and allowed to grow at room temperature. Growth curves to assess tolerance to nitrosative or oxidative stress were performed in LB broth at 37°C with shaking, starting from an overnight culture normalized to optical density at 600 nm (OD6_00_) = 0.05. At 2 hours post-inoculation, cultures were split into equal volumes and treated with 0.5 mM NOC-12, 1 mM H_2_O_2_, or left unperturbed. OD_600_ and CFU per mL measurements were taken every hour for 8 hours.

### RT-qPCR

RNA was extracted from day 11 colony biofilms or planktonic cultures using the RNeasy kit (Qiagen). RNA was DNase treated using Turbo DNase I (Invitrogen), and reverse transcribed using SuperScript III Reverse Transcriptase (Invitrogen). cDNA was amplified in an Applied Biosystems StepOne Plus Real-Time PCR machine using TaqMan MGB chemistry with the primers and probes listed in Table S1. All reactions were performed in triplicate with four different cDNA concentrations (100, 50, 25, or 12.5 ng per reaction). Relative fold difference in transcript abundance was determined using the ΔΔC_T_ method (42) with target transcripts normalized to *gyrB* abundance from a total of 3 – 4 biological replicates.

### Peptide nucleic acid fluorescence *in situ* hybridization (PNA-FISH)

Day 11 biofilms were flash frozen in Tissue-Tek O.C.T. compound (Electron Microscopy Sciences) and cryosectioned as described previously (5). The PNA-FISH hybridization protocol was adapted from Almeida *et al* (43). Biofilm cryosections were fixed in 4% paraformaldehyde (PFA) for 30 minutes at room temperature, then dehydrated for 10 minutes in 50% ethanol. After dehydration, 100 μL of hybridization solution (see below for details) was applied to the slides. All hybridizations were performed at 60°C for 30 minutes. Next, slides were submerged in pre-warmed wash solution for 30 minutes, mounted using ProLong Diamond (ThermoFisher), and imaged using a Zeiss 710 confocal laser scanning microscope (CLSM). For planktonic cells, 1 mL of culture was sedimented, fixed in 4% PFA, resuspended in 50% ethanol, incubated at −20°C for 30 minutes, and resuspended in 100 μL hybridization solution. After hybridization, cells were pelleted, resuspended in 500 μL pre-warmed wash solution, and incubated at 60°C for 30 minutes. Finally, cells were pelleted and resuspended in 100 μL sterile water before being applied to microscope slides for imaging. Wash solution contained 5 mM Tris-HCl pH 7.4, 15 mM NaCl, and 1% Triton X-100. Hybridization solution contained 10% w/v dextran sulfate, 30% formamide, 50 mM Tris-HCl pH 7.4, 10 mM NaCl, 5 mM EDTA, 0.1% Triton X-100, and 200 nM of each PNA-FISH probe. Probe sequences were based on the probes used for qPCR (efficiency of hybridization: *rrsH:* 81; *cydA*: 107; *cyoA:* 115; *appC*: 73) and were synthesized by PNA Bio (Newbury Park, CA).

### ATP measurements

ATP was quantified from mid-log (4 hours after subculture) planktonic cultures using Cell-Glo Titer kit (Promega). Cultures were normalized to OD_600_ = 0.5, pelleted, and resuspended in PBS. 50 μL of bacterial suspension was mixed with an equal volume of Cell-Glo Titer reagent and incubated shaking at room temperature for 15 minutes. After incubation, luminescence was measured on a SpectraMax i3 plate reader (Molecular Devices). Luminescence was converted to concentration of ATP using a standard curve on the same plate.

### Extracellular matrix extraction

Extracellular matrix was extracted using established methods (20). Briefly, biofilms were grown on YESCA agar containing 25 μg/mL Congo Red. After 60 hours, biofilms were homogenized in cold 10 mM Tris-HCl pH 7.4 using an Omni Tissue Homogenizer (motor speed 9) five times for one minute per cycle. Next, the homogenate was centrifuged three times for 10 minutes at 5,000g to remove cells. The supernatant was spiked with NaCl (final concentration 170mM) and centrifuged for one hour at 13,000g to pellet the matrix. The ECM pellet was washed in 10 mM Tris-HCl pH 7.4 with 4% SDS and incubated at room temperature rocking overnight. Next, the suspended ECM was centrifuged at 13,000g for one hour, resuspended in cold 10 mM Tris-HCl pH 7.4, and centrifuged at 30,000g for 20 minutes. Pelleted ECM was resuspended in MQ water and flash frozen.

### Solid-state NMR Measurements

All NMR experiments were performed in an 89 mm bore 11.7T magnet using either an HCN Agilent probe with a DD2 console (Agilent Technologies) or a home-built four-frequency transmission line probe with a Varian console. Samples were spun at 7143 Hz in either 36 μL capacity 3.2 mm Zirconia rotors or thin-walled 5 mm outer diameter Zirconia rotors. The temperature was maintained at 5°C with an FTS chiller (FTS Thermal Products, SP Scientific, Warminster, PA) supplying nitrogen at −10°C. The field strength for ^13^C cross polarization was 50 kHz with a 10% ^1^H linear ramp centered at 57 kHz. The CPMAS recycle time was 2 s for all experiments. ^1^H decoupling was performed with continuous wave decoupling. ^13^C chemical shifts were referenced to tetramethylsilane as 0 ppm using a solid adamantine sample at 38.5 ppm. The 15.6 mg wild-type ^13^C CPMAS spectrum was the result of 32,768 scans and the 8.3 mg mutant spectrum was the result of 100,000 scans. NMR spectra were processed with 80 Hz line broadening.

### SDS-PAGE gels

A portion of the lyophilized ECM sample used for solid-state NMR analysis was resuspended in 98% formic acid and vacuum centrifuged. The samples were then reconstituted in SDS-PAGE sample buffer containing 8M urea and 50 mM DTT and further diluted to desired concentrations. All samples were centrifuged briefly at 10,000g to remove any insoluble material and used for electrophoresis. The gels were stained with instant blue and de-stained in water.

### Immunofluorescence

Immunofluorescence targeting CsgA, the major curli subunit, was performed as previously described (10). Biofilm cryosections were fixed in 4% PFA for 30 minutes at room temperature and blocked overnight in 5% BSA at 4°C. Sections were washed in PBS, incubated with rabbit α-CsgA antibodies (GenScript) (1:1000) at room temperature for 1 hour, washed in PBS, and incubated with AlexaFluor 647 goat α-rabbit IgG (ThermoFisher) (1:1000) at room temperature for 1 hour. Slides were counterstained with SYTO 9 and imaged using CLSM.

### Murine infections

Murine infections were performed as described previously (44). In brief, UTI89 and each mutant strain were inoculated individually into 5 mL LB medium and grown shaking at 37°C for 4 hours. Next, this culture was diluted 1:1000 into 10 mL fresh media and grown statically at 37°C for 24 hours. After 24 hours, this culture was diluted 1:1000 into 10 mL fresh media and grown for another 24 hours at 37°C statically. Next, 7 – 8 week old C3H/HeN female mice were transurethrally inoculated with 50 μL PBS containing 10^7^ CFU bacteria. Mice were sacrificed at 24 hours post infection after which bladders were removed and homogenized for CFU enumeration. All animal studies were approved by the Vanderbilt University Medical Center Institutional Animal Care and Use Committee (IACUC) (protocol number M/12/191) and carried out in accordance with all recommendations in the Guide for the Care and Use of Laboratory Animals of the National Institutes of Health and the IACUC.

### Statistical analysis

All statistical analyses were performed in GraphPad Prism using the most appropriate test. Details of test used, error bars, and statistical significance cutoffs are presented in figure legends.

## Acknowledgments

We thank Dr. Jonathan Schmitz, Dr. Gerald Van Horn, Dr. Mariana X. Byndloss, and members of the Hadjifrangiskou laboratory for critical evaluation of the manuscript and helpful discussions. This work was supported by the following NIH grants: R01 AI107052 (MH), DiaComp DK076169 (MH), T32 GM007347 (CJB), K08 DK106472 (DBC) and the Vanderbilt University Medical Center Pediatric Urology Research Fund. LC acknowledges support from the National Science Foundation CAREER Award 1453247. SHW is a recipient of a NSF Predoctoral Fellowship.

## Author Contributions

CJB, ARE, and MH conceived the study, performed the experiments, analyzed the data, and composed the manuscript. SAR performed the nitrosative and oxidative stress assays. MJF and MMH constructed deletion mutants and complementation constructs under the supervision of CJB and ARE. JRB assisted with imaging data acquisition. SD and DBC performed the animal experiments along with CJB and ARE, assisted in data analysis, and edited the manuscript. SHW and LC performed the extracellular matrix component measurements and edited the manuscript.

---

**Figure S1:**
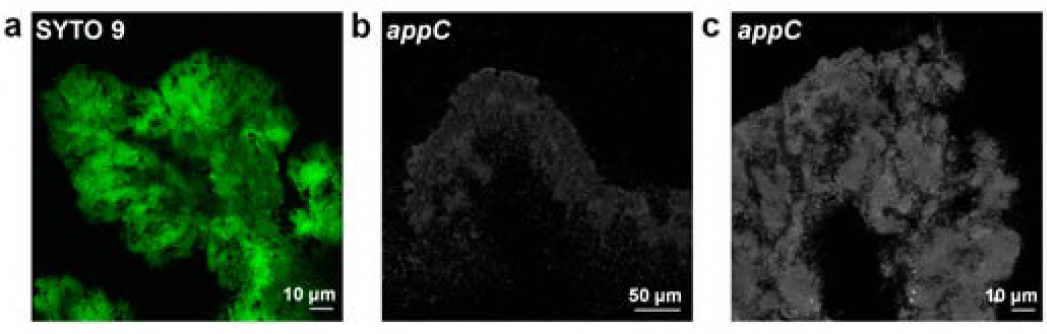
Localization of *appC* transcript in biofilm cryosections. **a**, Representative image of SYTO 9 stained biofilm cryosection at 20x magnification. **b – c**, Representative images of biofilm cryosections stained with an *appC* PNA-FISH probe at 20x (**b**) and 63x (**c**) magnification.

**Figure S2:**
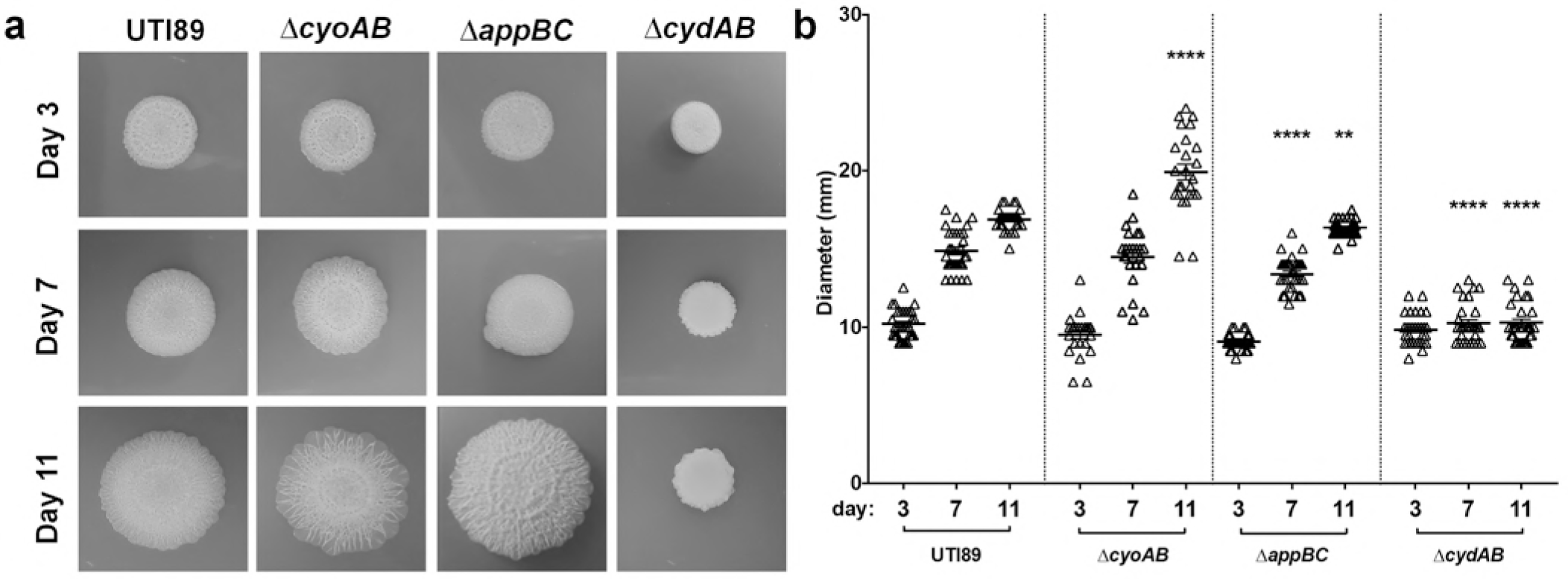
Temporal development of colony biofilms by UTI89 and isogenic cytochrome oxidase mutants. **a**, Representative images of UTI89 and cytochrome oxidase mutant colony biofilms grown on YESCA agar taken on day 3, 7, and 11 of growth. **b**, Graph depicting colony biofilm diameter at day 3, 7, and 11 of growth. Each triangle represents an individual colony biofilm. Data is representative of at least 30 biological replicates. Statistical analysis was performed in GraphPad Prism using Welch’s t test. * = p < 0.05, ** = p < 0.01, *** = p< 0.001, **** = p < 0.0001.

**Figure S3:**
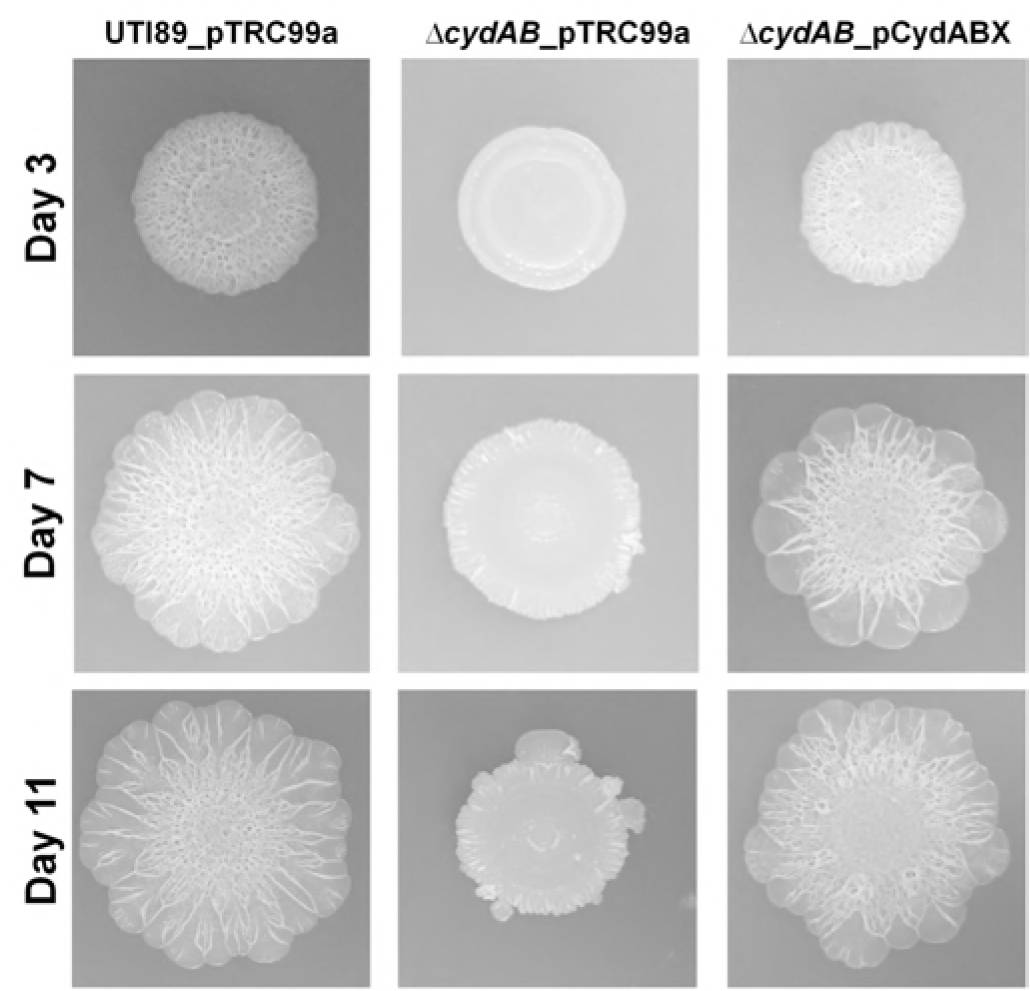
Extrachromosomal complementation of *ΔcydAB* rescues biofilm defects. Representative images on UTI89_pTRC99a, *ΔcydAB*_pTRC99a, and complemented *ΔcydAB*_pCydABX under the control of a native promoter. Images were taken of colony biofilms grown on YESCA agar at days 3, 7, and 11 of growth. Images are representative of at least 5 biological replicates.

**Supplemental Movies S1-4: Cytochrome *bd* influences matrix hydrophobicity and resistance to penetration by aqueous solutions.** Videos of colored water applied adjacent to a day 11 colony biofilm to assess the degree to which colony biofilms uptake aqueous solutions. **Movie S1**, UTI89; **Movie S2**, *ΔcyoAB;* **Movie S3**, *ΔappBC;* **Movie S4**, *ΔcydAB.* Videos are representative of at least 5 biological replicates.

**Figure S4:**
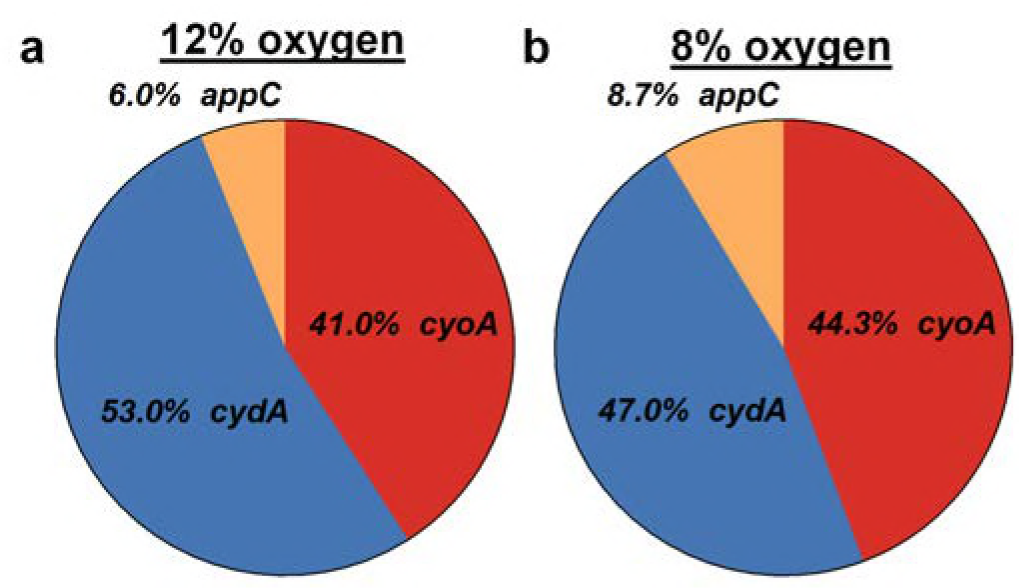
Ambient oxygen concentration influences cytochrome oxidase transcript abundance. Pie charts depicting relative abundance of *cydA, cyoA*, and *appC* transcripts detected using RT-qPCR in planktonic cultures grown at 12% (**a**) or 8% oxygen (**b**). Data is representative of three biological replicates.

**Figure S5:**
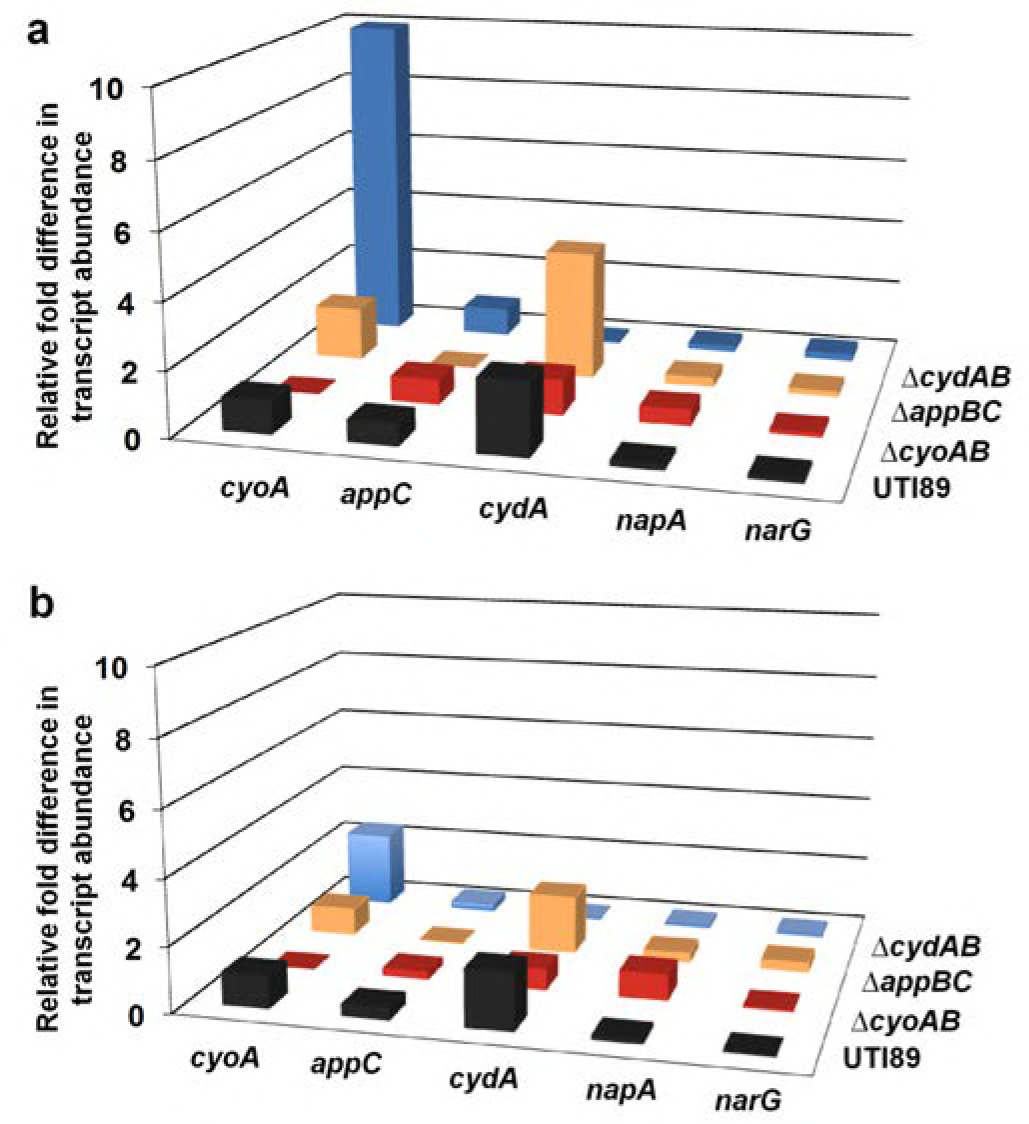
Expression of respiratory complexes in colony biofilms of cytochrome oxidase deletion mutants. RT-qPCR data depicting relative fold difference in respiratory transcript abundance in the center (**a**) and periphery (**b**) of day 11 colony biofilms across cytochrome oxidase mutant strains. In UTI89 (black), data is presented as relative fold difference in transcript abundance as compared to *cyoA* abundance. In each mutant strain, data is presented as relative fold difference in transcript abundance as compared to transcript abundance in UTI89.

**Table S1:**
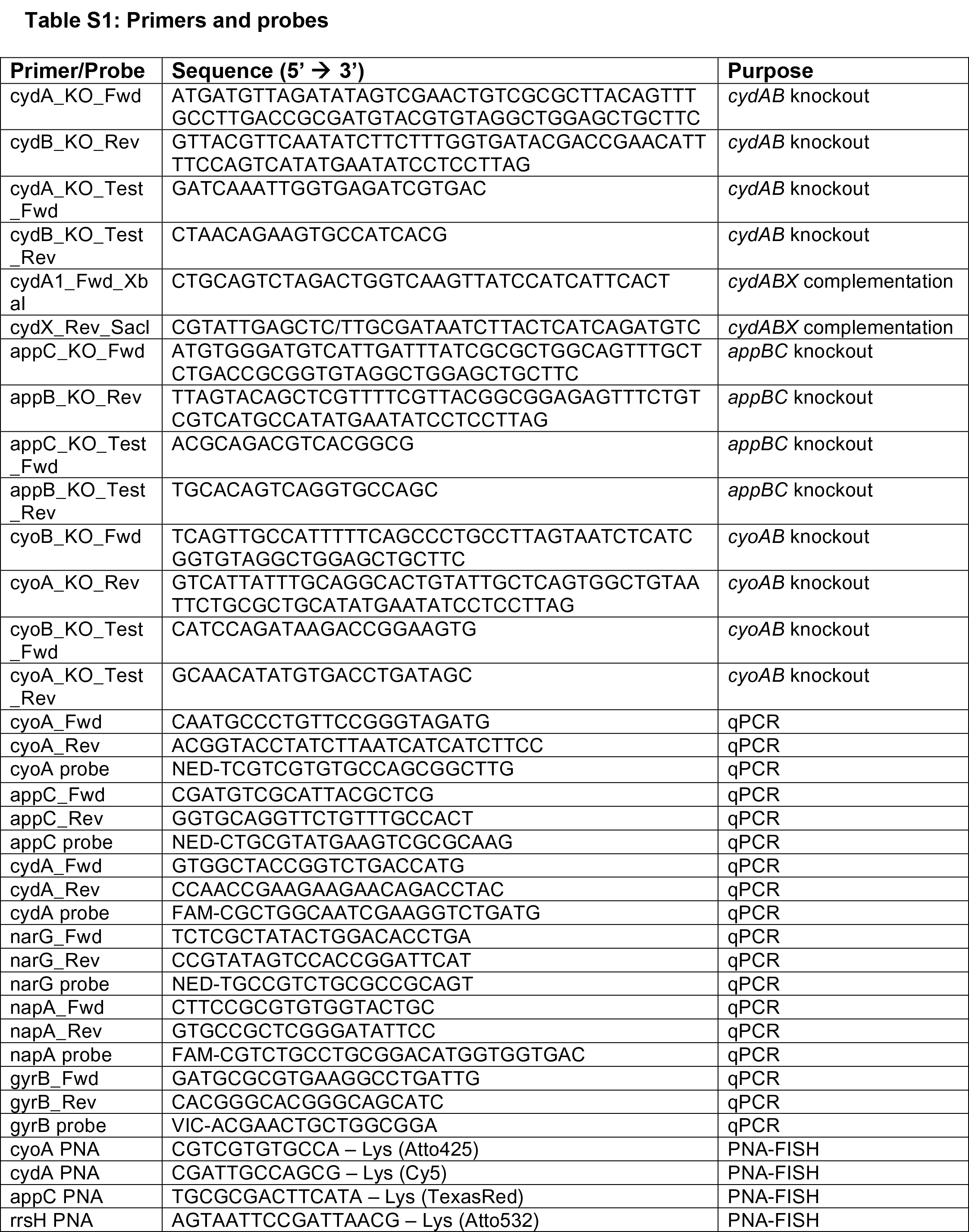
Primers and probes

